# Structural similarities of molecules selectively binding the *prfA* thermosensor RNA

**DOI:** 10.64898/2026.03.11.711090

**Authors:** Daniel Scheller, Rabindra Das, Erik Chorell, Jörgen Johansson

## Abstract

In light of the “silent” AMR pandemic, new avenues to combat pathogenic bacteria are needed. In this work, we screened a large molecule library (n=35 684 unique compounds) with the aim of identifying molecules being able to bind and block translation of the *prfA*-thermosensor transcript in the bacterial pathogen *Listeria monocytogenes*. Using a thiazole-orange displacement approach, 468 (∼1.3% of all molecules) showed the ability to reduce fluorescence. After dose response testing, 32 compounds remained promising and eight of them showed sufficient purity and availability to be further validated. Interestingly, four compounds, being structurally very similar, showed specificity for *prfA* at a varying degree. All four compounds carried 3 aromatic rings with one connecting amine between two of the rings and an amide linking an aliphatic amine side chain. The most selective compounds, M5, showed a K_d_ of ∼0.8 µM for the *prfA* RNA at 35°C. However, none of the eight most efficient compounds were able to inhibit *prfA* translation *in vitro*, suggesting that the molecules are able to bind but not affect the stability of the overall structure. Through this work, we have been able to identify a set of molecules, able to bind the *prfA* thermosensor RNA selectively, but without affecting translation. These molecules could constitute an important scaffold for further drug development.

## INTRODUCTION

It has been estimated that 1.27 million deaths in 2019 could be directly attributed to bacteria being resistant to one or several antibiotics (1). New types of antibacterial agents are thus required, not only for treating bacterial infections, but also to prevent infections from occurring (e.g., during complicated surgery and cancer treatments).

Several studies have highlighted the regulatory roles of secondary and tertiary RNA structures in various organisms, including bacteria (2). When being synthesized, the nascent RNA starts folding immediately from a primary unstructured state. One single-molecule transcript can adopt several distinct alternate secondary and tertiary structures depending on the free energy landscape of the RNA (3). The folding of particular structures can be affected by external physical factors (e.g. temperature) but also by macromolecules (e.g. metabolites, proteins or other RNAs). As a difference to proteins whose regulation is employed after their synthesis and folding, the regulatory mechanism exerted by regulatory RNA is continuous, starting immediately after transcriptional initiation and folding of the nascent RNA. In addition to classical Watson-Crick base-pairing, the chemical properties of the ribonucleotides (A, C, G, U), afford the formation of >10 non-canonical base-base interactions with different geometry as well as base-stacking, where bases stack on top of each other within nanoseconds (4). These unusual properties allow distinct RNA to be highly structured, and these RNAs will, like proteins, harbor specific pockets able to bind defined metabolites.

Targeting alternate RNA structures using small molecules has been suggested as a potential strategy to treat different diseases (5,6). For bacteria, it was shown that the small molecule Ribocil could bind an RNA regulatory region, block translation of the transcript and *Escherichia coli* infected mice treated with Ribocil reduced bacterial growth in the spleen (7). RNA-binding molecules differ from protein-binding molecules and typically interact with the structured RNA by two interfaces (4): These molecules typically contain several aromatic rings that can intercalate between stacked RNA bases, thereby strengthening the RNA-ligand interaction. In addition, RNA-binding molecules often contain amine groups that further stabilize the RNA-interaction when protonated at physiological pH *in vivo*, through ionic- and/or hydrogen-bonding with the RNA phosphate-backbone. Despite constituting a vast druggable space (e.g. 70% of the human genome is transcribed whereas only 2% is translated (8)), only a few clinical examples of RNA-binding drugs exist (9).

The food-born pathogen *Listeria monocytogenes* is the causative agent of listeriosis, a severe infection with a fatality of 20-30% in immunocompromised patients (10). *L. monocytogenes* is an intracellular pathogen, showing a distinct spatial and temporal expression of its virulence factors. Expression of *Listeria* virulence factors is tightly controlled at the transcriptional level by the main virulence-activating transcription factor PrfA (10). PrfA expression is controlled by an RNA thermosensor located in the 5’-UTR of the mRNA encoding PrfA (11). At low temperatures, the RNA-thermosensor inhibits ribosome binding, by occluding the ribosome binding site with its secondary structure, whereas it is opened to induce expression of virulence factors at host body temperature, due to melting of the RNA-structure. Importantly, a strain lacking PrfA expression is unable to cause disease (10). Thus, preventing re-structuring of the PrfA RNA thermosensor could be a possible strategy to decrease virulence factor expression and infectivity.

In this work, we set out to identify molecules able to 1) bind the *prfA* RNA thermosensor and 2) inhibit the opening of the *prfA* transcript to prevent translation of PrfA. Screening a molecular library of 35 684 unique molecules revealed four molecules showing a specificity for *prfA* at temperatures of infection. Although none of the identified molecules were able to inhibit *prfA* translation, they could constitute an important scaffold for developing new drugs targeting the *prfA* thermosensor RNA.

## MATERIALS AND METHODS

### Setup for HTS screen (35684 unique molecules)

#### Preparation of PrfA RNA

The PrfA RNA oligonucleotide was purchased from Integrated DNA Technologies. The oligonucleotide was dissolved in DEPC-treated RNase-free water to obtain a 100 µM stock solution and stored at −80 °C until use. Prior to the experiment, the stock solution was thawed and diluted to 100 nM in HEPES buffer (20 mM, pH 7.4) containing 150 mM NaCl, 5 mM MgCl₂, and 0.01 % Tween-20. The RNA solution was annealed by heating at 65 °C for 5 min followed by rapid cooling on ice for 30 min to allow proper folding.

#### Formation of the PrfA–Thiazole Orange (TO) Complex

A 10 mM stock solution of Thiazole Orange (TO) was prepared in DMSO and stored at −20 °C. Before the experiment, the TO stock was thawed and diluted to 100 µM using DEPC-treated water. The diluted TO was then added to the annealed RNA solution to achieve a final TO concentration of 400 nM. The mixture was incubated for 30 min to allow formation of the PrfA-TO complex. A sock plate with 125 µL PrfA-TO complex was prepared with matrix wellmate.

#### Ligand Displacement Assay and Screening

A total of 35,684 compounds (20 nL of 10 mM stock solution) from the compound collection at Chemical Biology Consortium Sweden were pre-spotted into 384-well assay plates (Thermo Fisher Scientific, Cat. No. 262260) using an Echo acoustic liquid handler. DMSO and RD310 were dispensed into columns 23 and 24 as negative and positive controls, respectively. Subsequently, 20 µL of the RNA-TO complex was dispensed into each well containing the pre-spotted compounds using a Biomek i5 Automated Workstation. This resulted in a final compound concentration of 10 µM. The plates were centrifuged using the integrated centrifuge system of the robotic platform and incubated for 30 min at room temperature prior to fluorescence measurement. Fluorescence was measured using a Clariostar microplate reader attached to the Biomek i5 Automated Workstation with excitation at 500±15 nm and emission at 533 ±15 nm. Compounds that produced a significant reduction in fluorescence relative to the negative control were considered potential hits, suggesting displacement of TO from the RNA or disruption of the RNA structure. Assay quality was evaluated using the Z′-factor (average 0.7) calculated from the control wells.

Percentage displacement was calculated using:

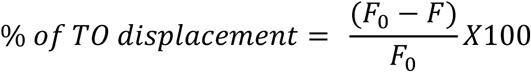

Where:

*F_0_*= fluorescence intensity of the RNA-TO complex without ligand

*F* = fluorescence intensity in the presence of ligand

A total of 468 compounds were selected based on their ability to reduce fluorescence intensity by at least 11.5% in the primary screening.

### Hit Confirmation

A three-point hit confirmation assay was performed on the 468 hits identified from the primary screen. The compounds were tested at three concentrations (5, 10, and 20 µM) in duplicate using the same assay conditions as in the primary screen. DMSO and RD310 (10 µM) were used as negative and positive controls, respectively.

Based on the results, 32 compounds were selected for further evaluation using an eleven-point full dose-response analysis in the TO-displacement assay.

### Setup for the dose-response experiment

An eleven-point dose-response experiment was performed on the 32 selected hits from hit confirmation using the same assay as in the primary screening. Compound transfer to assay plates was performed using an acoustic liquid handling system (Echo acoustic liquid handler) with the following final concentrations in triplicates: 100, 50, 25, 12.5, 6.25, 3.75, 2.5, 1.25, 0.625, 0.3125 and 0.125 µM. DMSO and RD310 (10 µM) were used as negative and positive controls, respectively. The percentage of TO displacement were evaluated using the same formula as in primary screening and dose-response curves were generated using GraphPad Prism 10 by plotting the percentage of TO displacement against compound concentration.

### Setup for the dose-response experiment

Prior to the experiment, the stock solutions of PrfA and HCoV were thawed and diluted to 200 nM in HEPES buffer (40 mM, pH 7.4) containing 300 mM NaCl, 10 mM MgCl₂, and 0.02 % Tween-20. The RNA solutions were annealed by heating at 65 °C for 5 min followed by rapid cooling on ice for 30 min to allow proper folding. TO solution (100 µM in DEPC-treated water) was added to the RNA solution (target concentration 800 nM) and incubated for 30 minutes to allow formation of the TO-RNA complex. A plate containing 10 µL of 20 µM (in DEPC treated water) selected hits in each well was prepared. Subsequently, 10 µL of the RNA-TO complex solution (either with PrfA or HCoV) was added to the wells and incubated for 30 minutes prior to fluorescence measurement using the same protocol. The TO displacement ability for the compounds were evaluated and bar diagram was plotted in excel.

### Microscale Thermophoresis (MST) experiments with *K*_d_ measurements

*prfA* or FSE RNA were Cy5 labelled at the 5′ end and folded in 20 mM HEPES and 150 mM NaCl, 5 mM MgCl_2_, and 0.01 % Tween 20 (pH 7.4) by heating at 65 °C for 5 min, followed by cooling down to room temperature. All MST experiments were performed in buffer containing 20 mM HEPES and 150 mM naCl, 5 mM MgCl2, and 0.01 % Tween 20. Labelled RNA concentration was held constant at 20 nM. For *K*d measurements, compound concentrations varied from 0.3 nM to 10 μM (16, 1:1 dilution steps). The samples were loaded into standard MST grade standard glass capillaries, and the MST experiment was performed using a Monolith NT.115 (Nano Temper, Germany). MST traces were plotted using OriginPro 8.5, and *K*_d_ values were generated by the NanoTemper analysis software.

### CD spectroscopy

Three μM amount of *prfA* or FSE RNA in 20 mM HEPES and 150 mM NaCl, 5 mM MgCl2, and 0.01 % Tween 20 (pH 7.4) by heating at 65 °C for 5 min, followed by cooling down to room temperature. A quartz cuvette with a path length of 1 mm was used for the measurements by a JASCO-720 spectropolarimeter (Jasco international Co. Ltd.). CD spectra of the RNA solutions were recorded either in the presence of DMSO or the hits (15 µM). Thermal melting curves for *prfA* and FSE RNA were recorded at 263 nm between 20 and 95 °C at a rate of 1 °C/min. Melting temperature (*T*_m_) is defined as the temperature at which 50% of the RNA structures are unfolded.

### Generation of DNA templates and *in vitro* transcription

The plasmid for *in vitro* transcription of the reporter fusion was generated by ligation of a PCR-amplified DNA fragment (respective primers T7-prfA-fw and prfA-EcoRI_rev), containing the T7 promoter, the RNAT and 15 bp of the coding region, into the EcoRI site of pZA23-eGFP. DNA Templates for *in vitro* transcription were generated by PCR amplification using T7-fw and eGFP-rev primers.

*In vitro* transcription was performed using the MEGAscript T7 Transcription Kit (Invitrogen) according to the manufacturer’s instructions with 200 ng of purified PCR products.

### *In vitro* translation

*In vitro* translation was carried out with PURExpress *in vitro* protein synthesis Kit (NEB) according to the manufacturer’s instruction. Briefly, 1 µg of *in vitro* transcribed RNA was used in a 10 µl standard reaction at either 30°C for 4h or 37°C for 2h in presence or absence of the tested compound. Total DMSO concentration in the reaction mixture was either 2 % or 3.35 %, depending on the compound concentration used. Translation products were further used for SDS-PAGE analysis and western blotting or fluorescence measurements.

### Fluorescence measurements of eGFP

5 µl of the *in vitro* translation product was mixed with 95 µl PBS and transferred to a black 96-well microtiter plate (Greiner bio-one). The fluorescence intensities were recorded using a multimode microplate reader (Tecan, Spark) at the excitation wavelength 485 nm and emission wavelength of 535 nm. PBS was used for background subtraction.

### SDS-PAGE and Western blot analysis

Of the *in vitro* translation product, 5 µl were mixed with 5 µl 1x SDS sample buffer (2% SDS, 0.1% bromophenol blue, 1% 2-mercaptoethanol, 25% glycerol, 50 mM Tris/HCl, pH 6.8) and boiled for 10 min at 95°C. Afterwards 10 µl of the supernatant was loaded and separated by SDS gel electrophoresis in 5% stacking and 12% separating gels. By semidry blotting, the proteins were transferred onto a nitrocellulose membrane (BioTrace NT, Pall Corporation) and a rabbit anti-eGFP antibody (Agrisera; AS21 4583) was used in a 1:2000 dilution, The next day, the membranes were washed 3 times with PBST and incubated with a goat anti-rabbit-HRP conjugate antibody (AS09 602, Agrisera) in a 1:10000 dilution. Chemiluminescence signals were detected by incubating membranes with Amersham ECL prime western blotting detection reagent (Cytiva) for 3 minutes.

Constructs were generated in a similar fashion as described in [15]. Briefly, a DNA template containing a T7 promoter, the *prfA*-thermosensor RNA, parts of the *prfA* coding region and *gfp* was synthesized by Eurofins Genomics into the vector. *In vitro* transcription was done with the MEGAscript T7 Transcription Kit (Invitrogen) according to the manufacturer’s instructions with 1 µg of *Sma*I digested pEX-constructs as template.

## RESULTS

### Primary screen reveals a subset of molecules able to bind the *prfA*-RNA

In an effort to identify molecules able to bind the *prfA* thermosensor RNA and inhibit translation of the PrfA protein, we undertook a thiazole orange (TO) displacement strategy. TO becomes highly fluorescent when binding RNA and molecules able to compete with TO for RNA binding reduce the observed fluorescence signal. A dose-response experiment was performed using 100 nM PrfA and varying concentrations of TO. The results indicated that 4 equivalents of TO produced stable signals. The *prfA*-RNA and TO (ratio of 1:4) was pre-incubated for 30 minutes at 25°C, before being exposed to each of the 35 684 molecules at 10 µM concentration in the primary screening set from Chemical Biology Consortium Sweden (CBCS). After incubation for an additional 30 minutes at 25°C the fluorescence readout was measured. As a positive control, we used RD310, a promiscuous nucleic acid binding molecule (12). 468 (∼1.3% hit-rate) of the molecules displayed a decreased fluorescence (at least 11.5% reduction – an arbitrarily selected cutoff) in comparison to the control and were characterized further (Figure 1A).

**Figure 1.**
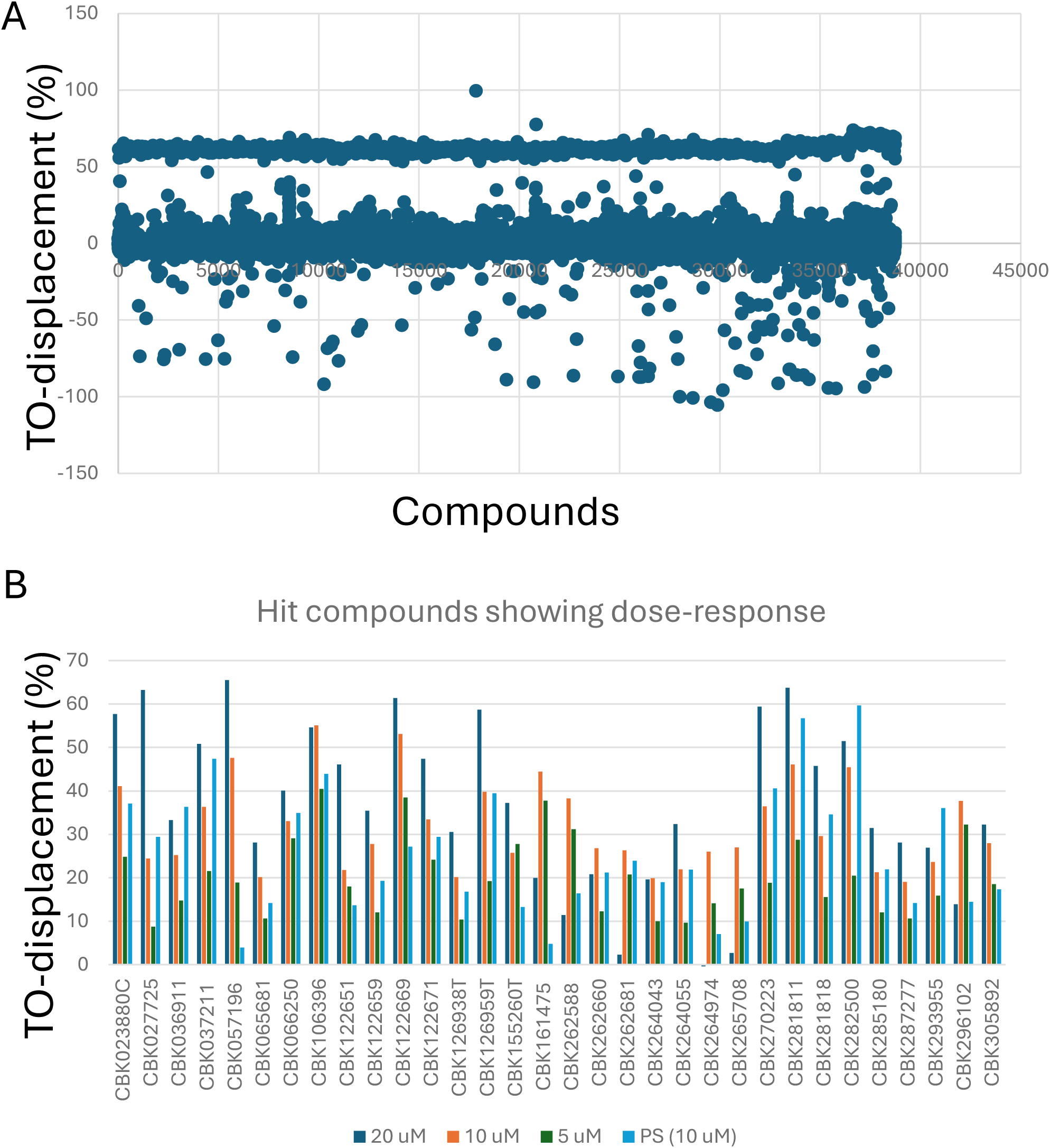
**A.** Primary screening approach by thiazole orange displacement assays to identify molecules able to bind the *prfA*-RNA thermosensor. 35 684 molecules were screened for their ability to displace thiazole orange (TO). TO and *prfA* RNA (4:1 ratio) were pre-incubated at 25°C for 30 minutes before being exposed to compounds (in 384 well microtiter plates) at 25°C for 30 minutes before fluorescence measurement **B.** 32 top-candidates selected by dose-response experiments of the 468 compounds identified from A. Dose-response experiments were performed at 5; 10 and 20 µM concentration, respectively. PS: primary screen.

### Dose response experiments further narrowed down the number of molecule candidates

We next performed hit-validation experiments at 5, 10, and 20 µM, respectively, of each of the 468 molecules identified in the primary screen. 32 molecules showed a clear dose response, with 10 molecules being able to displace at least 50% of the TO signal (Figure 1B). However, only 8 of the identified molecules were available at sufficient concentration and/or purity to allow further examination.

This unusually high prevalence of hits with insufficient purity suggests that the TO displacement assay may be particularly sensitive to assay-interfering species. In addition to genuine competition for RNA binding, reductions in fluorescence may arise from fluorescence quenching, compound aggregation, or other nonspecific interactions affecting the TO signal. However, control experiments, including TO-only and unrelated RNA controls, showed that the remaining candidates did not reduce fluorescence in the absence of the target RNA, supporting the interpretation that the prioritised compounds act through RNA-dependent interactions rather than nonspecific optical interference.

### The most promising molecules are structurally similar and bind the *prfA*-RNA selectively

Although binding *prfA* in a dose-dependent manner, it could be hypothesized that the molecules bind RNA non-specifically. To examine this, we took advantage of the Frameshifting Stimulating Element (FSE) RNA from the coronavirus HCoV-OC43. The FSE halts the elongating ribosome and in approximately 50% of the cases, the ribosome slips back one frame (-1 programmed frameshift) allowing continuous translation of > 4000 amino acids. In case of no frameshift, translation will shortly terminate at a stop-codon. Importantly, the FSE is highly structured, forming a stable 3D RNA structure (13). Examining binding of the eight most potent molecules revealed that two of them (C5 and E3) displayed a higher affinity for the FSE-RNA compared with the *prfA*-RNA (Figure 2A). While two other molecules (E5 and O3) showed an equal affinity for the FSE-RNA and *prfA*-RNA, four molecules (M5, K5, O5 and C7, showed stronger affinity for the *prfA*-RNA than for the FSE-RNA, with M5 showing the largest selectivity. Examining the structure of the four selective molecules revealed striking similarities, all had a central pyrazine ring flanked by two phenyl substituents, one directly linked and the other linked by an amine group (Figure 2B). In addition, each molecule possessed an amide linkage to an aliphatic amine side chain. This molecular architecture is characteristic for many RNA-binding molecules since the of aromatic groups can intercalate between stacked base pairs whereas the amines may form hydrogen bonding or electrostatic interactions with the phosphate groups in the RNA backbone (4).

**Figure 2.**
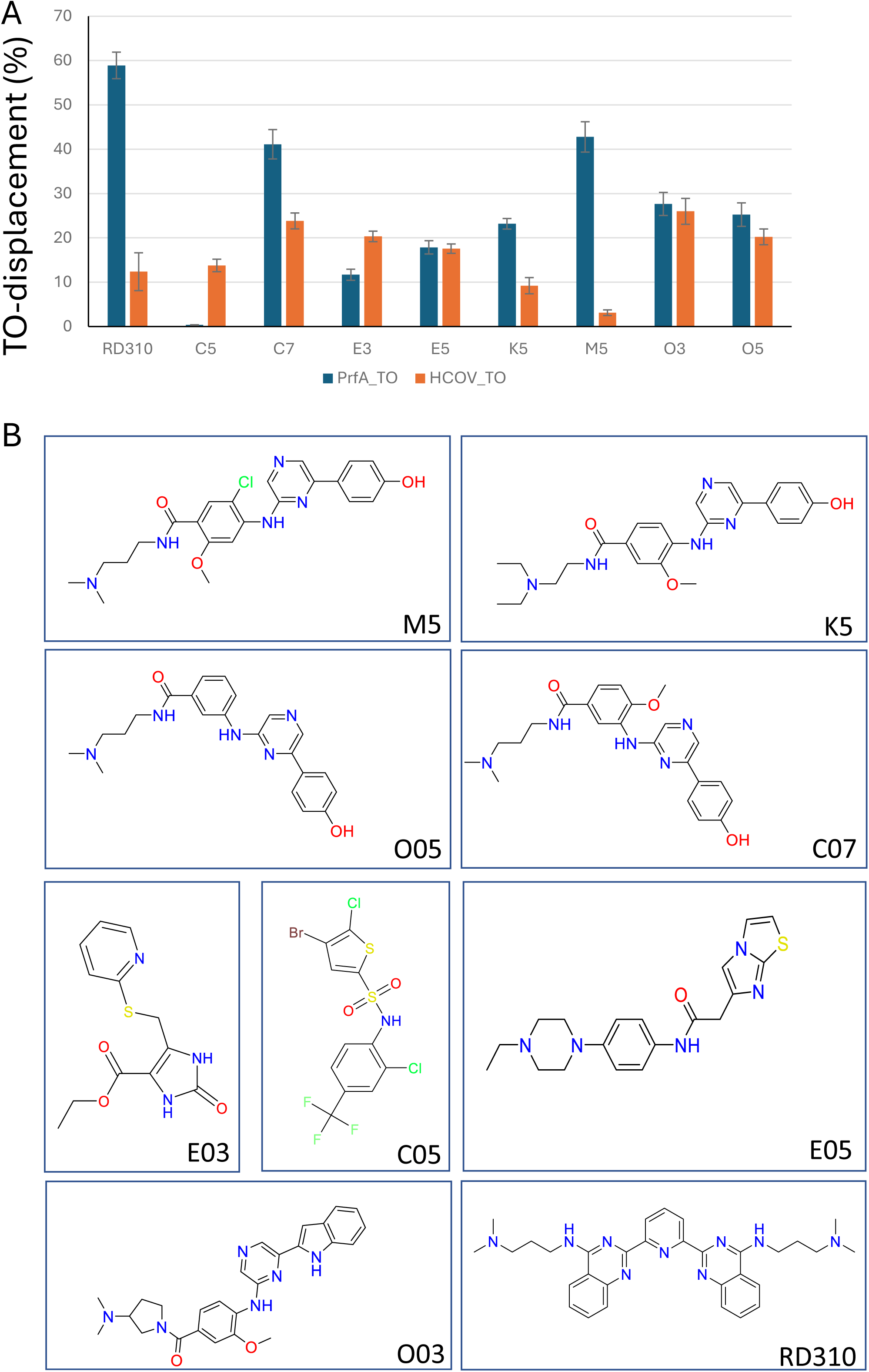
**A.** Most promising molecules bind selectively to the *prfA*-RNA. TO and *prfA* RNA or TO and HCoV RNA (4:1 ratio) were pre-incubated at 25°C for 30 minutes before being exposed to compounds. The samples were incubated at 25°C for 30 minutes before fluorescence measurement (TO: thiazole orange). **B.** Structure of the eight most potent molecules.

### The four most promising molecules bind the *prfA*-RNA at 35°C

To examine these molecules further, we performed Microscale thermophoresis (MST) measurements focusing on M5. Interestingly, we observed that M5 only could interact with the *prfA*-RNA at 35°C and not at 25°C, whereas RD310 could interact at both temperatures. None of the molecules could interact with the FSE-RNA at neither 25°C or 35°C (Figure 3A). Through MST, we calculated the *K*d value for M5, K5, O5 and C7, respectively (Figure 4B). Strikingly, the most selective molecule (M5) also showed the strongest affinity for the *prfA*-RNA (*K*_d_ ∼0.82 µM) whereas the other molecules showed a decreased affinity in direct correlation with the selectivity (Figure 3B). Finally, Circular Dichroism (CD) analysis was performed to examine whether the thermal unfolding (melting) of the *prfA*-RNA was affected by the different molecules. No large differences were observed for M5, K5, O5 and C7, whereas RD310 slightly stabilized the *prfA*-RNA structure resulting in an increased melting temperature (Figure 3C). None of the molecules (including RD310) were able to affect the melting temperature of the FSE-RNA (Figure 3D).

**Figure 3.**
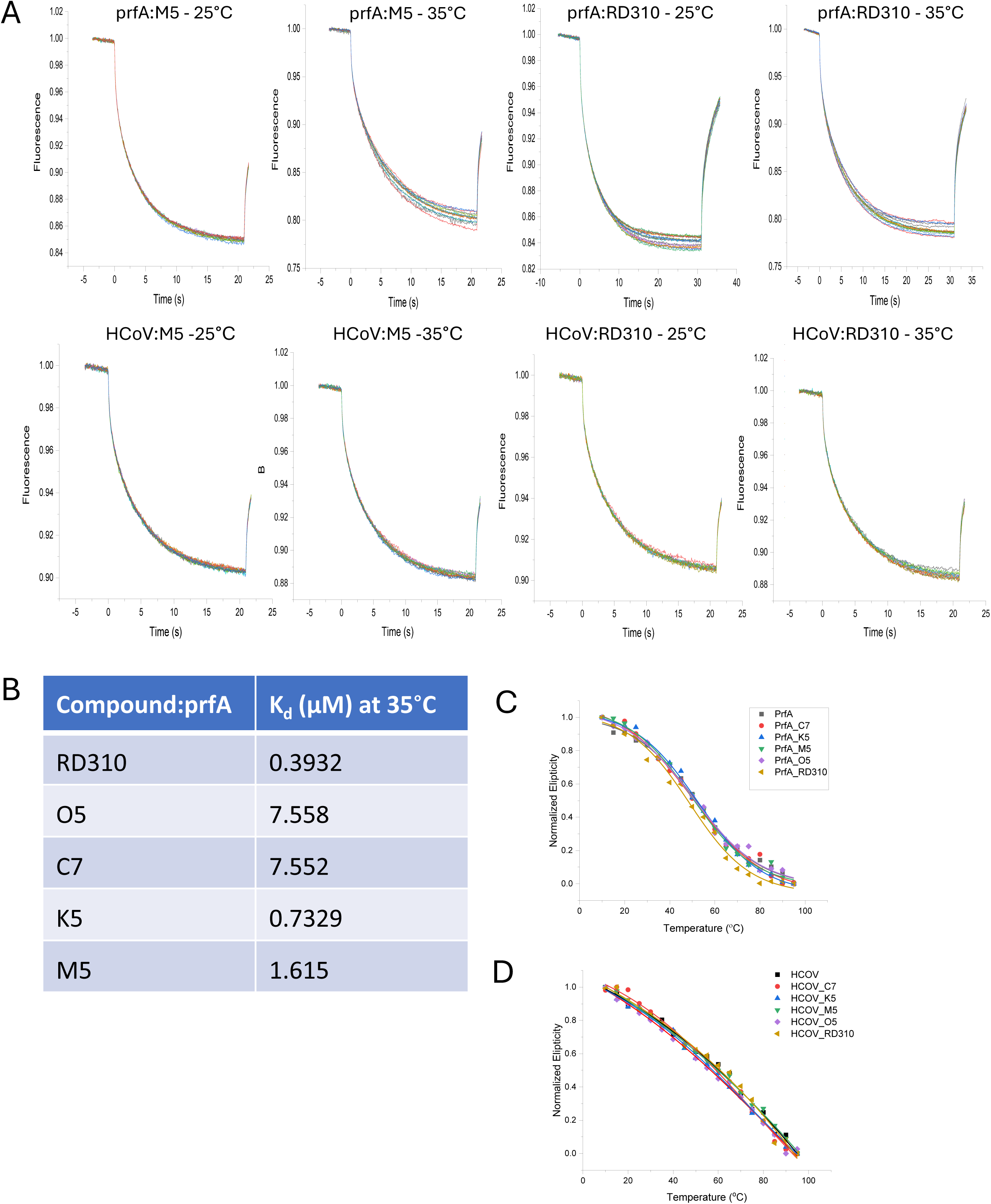
**A.** M5 bind selectively to the *prfA*-RNA at 35°C, but not at 25°C. *prfA* or HCoV (FSE) RNA (20 nM) were denatured and allowed to cool to room temperature. For *K*d measurements, compound concentrations varied from 0.3 nM to 10 μM (16 1:1 dilution steps) **B.** Table of *K*d measurements. **C** and **D.** Thermal melting curves for *prfA* (A) and FSE (B) RNA between 20 and 95 °C. CD spectra of the RNA solutions (3 μM) were recorded either in the presence of DMSO or the hits (15 µM).

**Figure 4.**
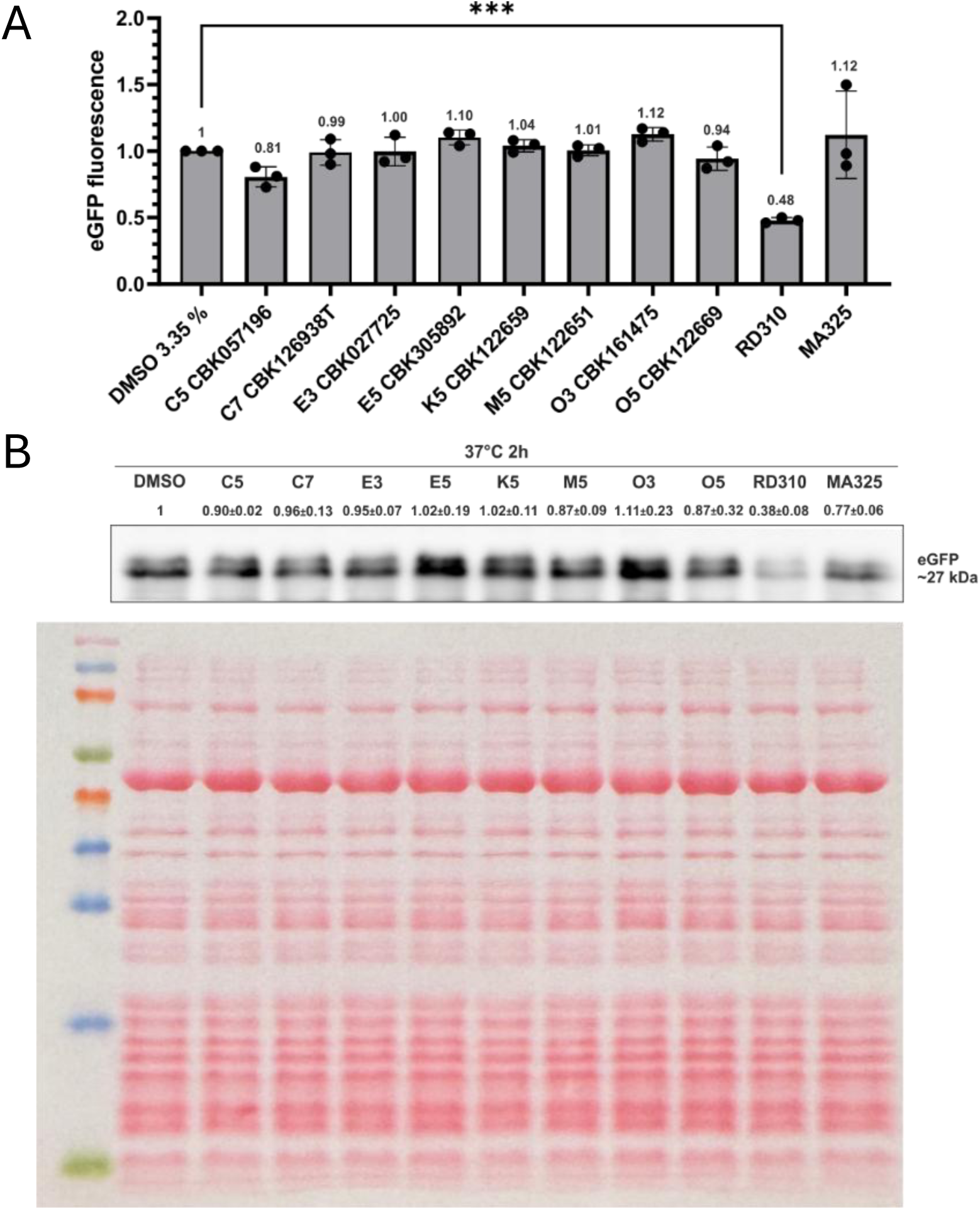
**A.** None of the eight most potent molecules can inhibit *prfA* translation at 37°C. Fluorescence intensities of *in vitro* translation products were measured at 485 nm excitation and 535 nm emission. Experiments were carried out three times. Mean and corresponding standard deviation are shown. All values have been normalized, relative to the control translation as 1. Fold differences to the DMSO control are indicated above the bars. **B.** *In vitro* translation of the PrfA-eGFP reporter by PURExpress *in vitro* translation kit in presence or absence of compounds. Experiments were carried out three times. Signals have been quantified using ImageJ and all values have been normalized, relative to the control translation as 1. Fold differences to the DMSO control are indicated as mean with the corresponding standard deviation. One representative Western blot is shown.

### The eight most potent molecules do not affect translation

Since we observed binding of the eight molecules to the *prfA*-RNA at varying affinity and selectivity, we were interested in examining whether they could affect translation *in vitro*. To examine this, we used an *in vitro* translation assay, measuring production of eGFP. In contrast to the positive control RD310, none of the eight molecules were able to significantly affect *prfA* translation (Figure 4A). To exclude the possibility that the observed effects were due to indirect fluorescence quenching, we also measured the amount of PrfA-eGFP protein being produced by western blot analysis. The results closely mirrored the data obtained through fluorescence measurements (Figure 4B).

## DISCUSSION

In this work, we have examined whether the *prfA*-RNA thermosensor could be a druggable target. Screening of a large molecule library containing 35 684 unique molecules identified four compounds showing selectivity for *prfA*, with one, M5, having a *K*_d_ of ∼0.8 µM. Further examinations, like DMS-MapSeq assays, could reveal the exact binding site of the molecules on the *prfA*-RNA thermosensor.

Interestingly, the four most selective molecules were structurally very similar, each containing a central pyrazine ring flanked by two phenyl rings, one directly attached and the other connected via an amine linker, and an additional amide linkage to an aliphatic amine side chain. (Figure 2B). Aromatic rings and amine-linkers have been suggested to distinguish RNA-binding molecules from protein-binding molecules.

The aromatic rings may intercalate between stacked base-pairs, while the aliphatic amines can form hydrogen bonding or electrostatic interactions with the phosphate groups in the RNA-backbone. Notably, all four of the most selective molecules identified in this screen share these structural characteristics. Furthermore, it is interesting to note that subtle differences in substituents at specific positions greatly influence both binding affinity and selectivity.

Unfortunately, none of the molecules being able to selectively bind the *prfA*-RNA, could inhibit translation *in vitro*. The reasons for this are not clear. One possible explanation is that compound binding alone is insufficient to prevent the structural rearrangements required for translational initiation. Small molecules that associate with structured RNA frequently stabilize local base stacking or conformations rather than locking the RNA in a single global inhibitory structure. Consequently, ligand binding may not prevent, and could potentially even facilitate, the unfolding events required for ribosome access.

Another possibility is that the molecules bind to regions of the RNA that do not directly control the translational initiation. In such cases, selective binding would not necessarily translate into functional inhibition.

Finally, it is also possible that the affinity of the compounds is insufficient to compete with ribosome binding or with the structural rearrangements occurring during translational initiation. Together, these possibilities suggest that selective RNA binding alone may not be sufficient to inhibit translation and that effective inhibitors may instead need to stabilize a specific inhibitory RNA conformation. However, selective RNA-binding molecules may still prove highly valuable in alternative RNA-targeting strategies. For example, the selectivity of these compounds, particularly M5, could prove valuable in a bifunctional small molecule designed to bind the target RNA while simultaneously recruiting or inducing RNA cleavage (14). RiboTACs (ribonuclease targeting chimeras) are RNA-binding molecules tagged with an RNA degradation extension that recruits an endogenous RNase (15). Other variants are RNA-binding molecules conjugated with bleomycin or imidazole. In these examples, bleomycin and imidazole carry out the “RNase”-activity themselves (16,17). Creating a variant of M5 conjugated with one or another of the RNA-degradation features could thus be an affordable path forward.

## ACKNOWLEDGEMENTS

J.J. was funded by the Swedish Research Council grant #2023-02679, Umeå University, the Stiftelsen Olle Engkvist Byggmästare, Vinnova grant 2019-05491 and the Erling-Persson Foundation.

## Notes

### Competing Interest Statement

The authors have declared no competing interest.

